# The impact of genome-wide association studies on biomedical research publications

**DOI:** 10.1101/106773

**Authors:** Travis J. Struck, Brian K. Mannakee, Ryan N. Gutenkunst

## Abstract

The past decade has seen major investment in genome-wide association studies (GWAS), with the goal of identifying and motivating research on novel genes involved in complex human disease. To assess whether this goal is being met, we quantified the effect of GWAS on the overall distribution of biomedical research publications and on the subsequent publication history of genes newly associated with complex disease. We found that the historical skew of publications toward genes involved in Mendelian disease has not changed since the advent of GWAS. Genes newly implicated by GWAS in complex disease do experience additional publications compared to control genes, and they are more likely to become exceptionally studied. But the magnitude of both effects has declined dramatically over the past decade. Our results suggest that reforms to encourage follow-up studies may be needed for GWAS to most successfully guide biomedical research toward the molecular mechanisms underlying complex human disease.

**Author summary:** Over the past decade, thousands of genome-wide association studies (GWAS) have been performed to link genetic variation with complex human disease. A major goal of such studies is to identify novel disease genes, so they can be further studied. We tested whether this goal is being met, by studying patterns of scientific research publications on human genes. We found that publications are still concentrated on genes involved in simple Mendelian disease, even after the advent of GWAS. Compared to other genes, disease genes discovered by GWAS do experience additional publications, but that effect has declined dramatically since GWAS were first performed. Our results suggest that the ability of GWAS to stimulate research into novel disease genes is declining. To realize the full potential of GWAS to reveal the molecular mechanisms driving human disease, this decline and the reasons for it must be understood, so that it can be reversed.

## Introduction

Since the first successful genome-wide association studies (GWAS) were published over a decade ago [1–3], thousands have been performed [4]. These studies have identified tens of thousands of statistical associations between genetic variants and human diseases [4]. More broadly, they have provided insight into the genetic architecture of complex human disease, showing that most common diseases are polygenic and that most common variants only slightly affect disease risk [5]. The large investment in GWAS has been criticized [6], perhaps because initial hopes for quick clinical impact were overenthusiastic [7]. The average time from basic science discovery to clinical practice is 17 years [8], so it unsurprising that few GWAS results directly affect patients yet. But direct clinical impact is not the only goal of GWAS.

A major goal of GWAS is to identify novel genes involved in complex disease and steer research toward them [9–11]. For example, an early GWAS unexpectedly found variation in Complement Factor H to be strongly associated with macular degeneration [1], spurring the development of complement-based therapeutics [12]. Similarly, associations between variation in the Interleukin 23 Receptor and Crohn’s disease [13] and psoriasis [14], motivated the development of several treatments that are now in clinical trials [15]. In both of these classic examples, going from association to therapy demanded substantial follow-up research.

To assess the impact of GWAS on biomedical research, we focused on scientific publications. Published GWAS are themselves often highly cited, for example [3, 13, 16]. A systematic comparison also found that GWAS are more highly cited than comparable candidate gene studies [17]. But a paper that cites a GWAS does not necessarily follow-up on the associations reported by that GWAS. To quantify how much follow-up research is motivated by GWAS, we focused on the subsequent publication record of newly associated genes.

The distribution of biomedical research publications is highly unequal among human genes (Fig. 1A) [18]. Much of this inequality stems from historical momentum, driven by the availability of prior functional information [19] or research tools [20].

**Fig 1.**
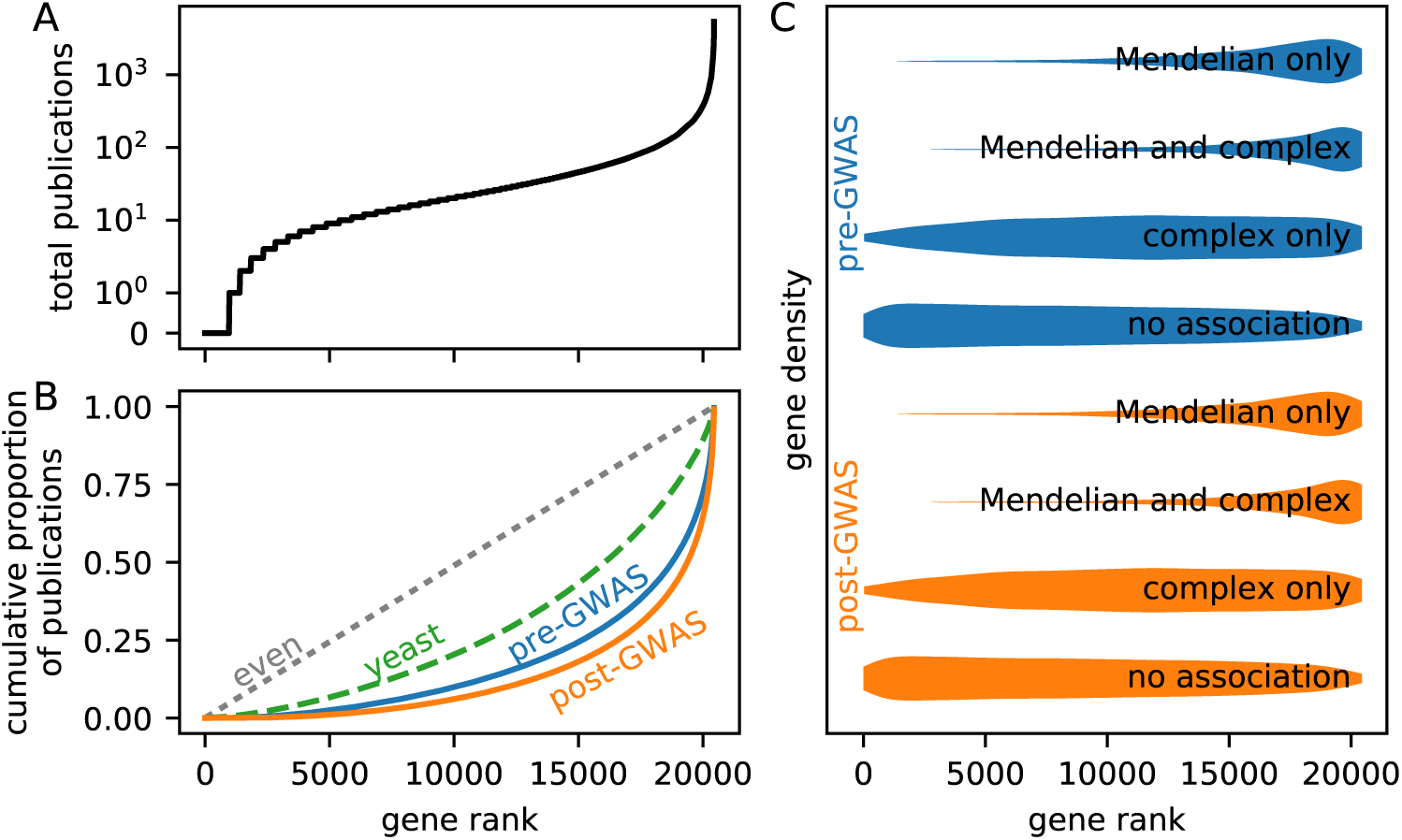
Biomedical scientific publications are highly unequally distributed and strongly skewed toward genes involved in Mendelian disease, even after the advent of GWAS. A: The distribution of publications among human genes is highly uneven. Plotted is the number of publications per gene, with genes sorted by number of publications. A few genes are the subject of thousands of publications each, whereas thousands of genes are the subject of fewer than ten publications each. B: The distribution of publications among human genes is more uneven in the post-GWAS era (2005 and later) than in the pre-GWAS era (before 2005). Shown in this Gini plot are the cumulative proportions of publications in each category, versus gene rank. The further the curve is from the diagonal, the more uneven the distribution. For comparison, the distribution of publications among yeast genes is shown, with the yeast x-axis stretched to match the number of human genes. C: Highly-studied genes tend to be involved in Mendelian disease. Plotted is the density of genes versus publication rank for genes of each possible type of disease association and for both the pre- and post-GWAS eras. In both eras, genes involved in Mendelian diseases are strongly enriched toward high publication ranks. By contrast, many genes involved only in complex disease rank low in terms of publications.

Consequently, many potentially medically important genes may be understudied [21]. Because GWAS are largely unbiased by previous knowledge about genes [22], they provide an opportunity for understudied genes to be brought to the scientific forefront.

We evaluated the effect of GWAS on the biomedical research literature in three ways. At a broad scale, we tested whether the distribution of publications among human genes has changed since the advent of GWAS. At a narrower scale, we quantified the effect of being newly associated with complex disease on the subsequent publication histories of human genes. Lastly, we identified outlier genes with exceptional publication activity and tested whether GWAS might play a role in motivating such activity. Overall, we find that genes newly associated with complex disease do experience notable increases in publication activity, but this effect has declined dramatically over the past decade.

## Results

We measured research output on genes using scientific publications, as collected in the NCBI Gene database [23]. We prefer this manually curated database to automatic text mining, because text mining may introduce false positives when a gene is mentioned in passing. We used the Online Mendelian Inheritance in Man (OMIM) database [24] and the EBI-NCBI GWAS catalog [4] to classify genes into those associated with no disease, Mendelian disease, complex disease, or both.

### Broad patterns of publications on human genes

As expected [18], we found that the distribution of publications among human genes was highly uneven. A small number of genes were the subject of many thousands of publications, while a large number of genes were the subject of only a few (Fig. 1A).

To quantify the unevenness of publications among genes, we used the Gini coefficient, which ranges from 0 (perfectly even distribution) to 1 (perfectly uneven). The Gini coefficient is calculated from the cumulative distribution of publications versus gene rank (Fig. 1B). Notably, the inequality of publications among human genes is larger in the post-GWAS era than in the pre-GWAS era (Gini coefficient 0.73 vs 0.65; Fig 1B). It is not inevitable that the distribution of publications should be so unequal; the Gini coefficient of publications among yeast genes is much lower at 0.43 (Fig. 1B).

The ultimate goal of most biomedical research is to improve human health, so the distribution of publications is expected to be skewed toward genes involved in human disease. In the pre-GWAS era, genes associated with Mendelian disease were, almost without exception, among the most highly studied human genes (Fig. 1C). By contrast, many genes that would later be associated with complex disease were among the least studied human genes (Fig. 1C). The advent of GWAS led to the discovery of many genes associated with complex human disease. The focus of biomedical publications on Mendelian disease genes, however, remains strong in the post-GWAS era (Fig. 1C). In particular, many genes associated with complex disease remain among the least studied genes in the human genome (Fig. 1C).

The advent of GWAS has thus not reduced the startling inequality of scientific publications among human genes (Fig. 1B) nor qualitatively changed the skew of those publications toward genes involved in Mendelian, but not complex, disease (Fig. 1C). But how do GWAS affect subsequent publications on individual genes that are newly associated with complex disease?

### Subsequent publications on individual genes

To quantify the immediate effect of GWAS on research into individual newly associated genes, we focused on the calendar year a gene was associated with complex disease through GWAS and the following two years. For each new GWAS gene, we compared publications over this period with a control non-GWAS gene chosen to have as similar a prior publication history as possible. The variance in an associated gene’s publications is strongly correlated with the number of publications on that gene in the prior three years (Fig. 2A). Normalizing the excess in publications relative to the control gene by the square root of the number of recent publications normalizes the variance (Fig. 2B), consistent with a Poisson model for publication output [25]. The normalized excess in publications for a GWAS gene is slightly but significantly shifted (Fig. 2C; one-sample *t*-test, *p* ∼5 ×10^*-*34^, *N* = 2, 442). The mean normalized excess is 1.24 units, corresponding to a mean excess of 2.95 publications over the three years following association.

**Fig 2.**
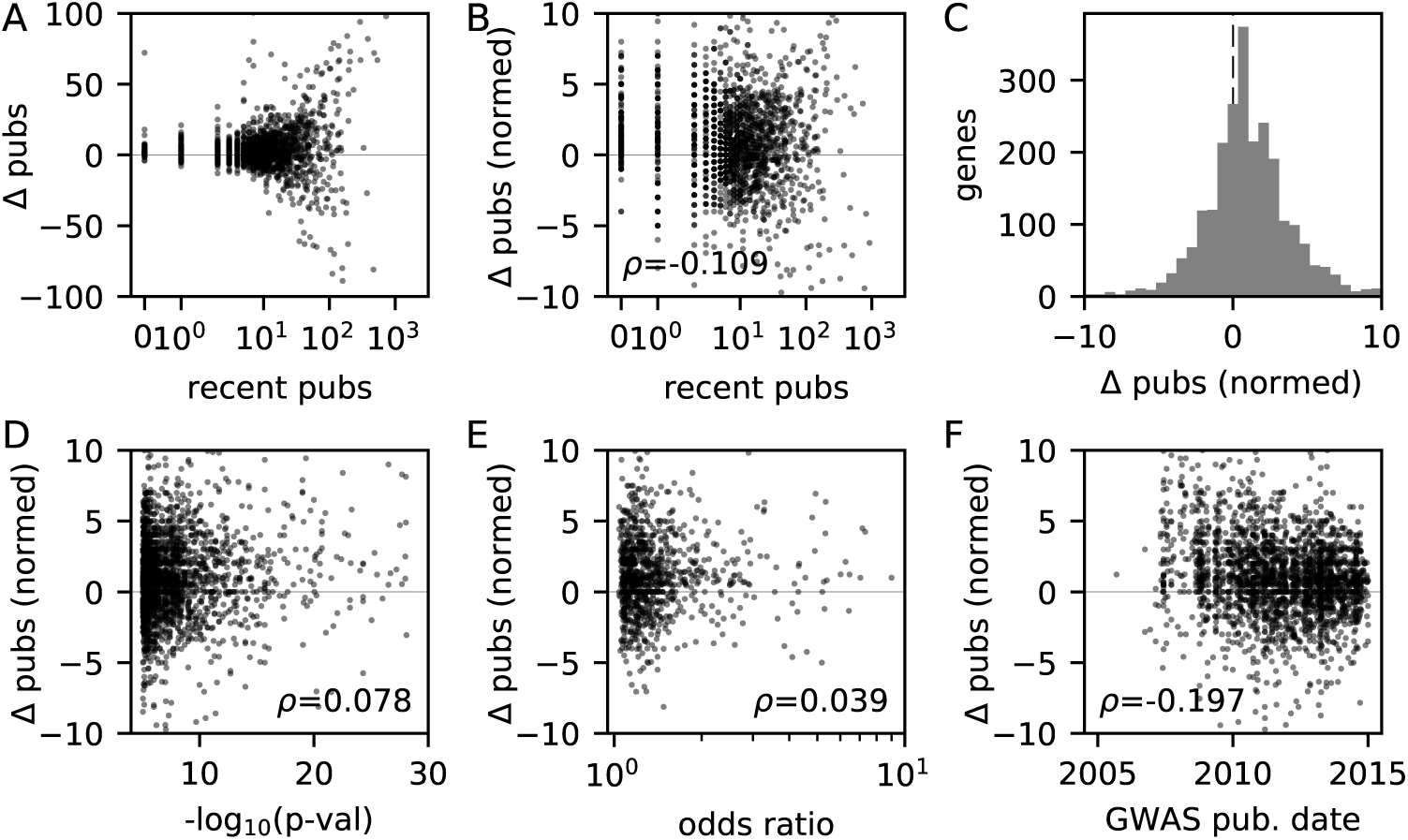
Effect on subsequent publications for genes newly associated with complex disease via GWAS. To quantify the short-term effects of GWAS association, we considered the publication excess of each newly associated gene compared with its control gene. A: The variance of the publication excess is strongly correlated with the associated gene’s number of recent publications. B: Normalizing the publication excess by the square root of the number of recent publications equalizes the variance. It also reveals a trend for the normalized effect of GWAS association to be smaller for more heavily studied genes. C: The distribution of normalized publication excess is shifted toward positive values, indicating a positive effect of GWAS association on subsequent publications. D: The normalized publication excess for a newly associated gene is weakly correlated with the p-value of the association. E: It is not statistically significantly correlated with the effect size of the association, as quantified by the odds ratio. F: The normalized publication excess is negatively correlated with the publication date of the association. More recently associated genes experience a smaller increase in subsequent publications.

What factors determine how large an effect a GWAS will have on an associated gene’s subsequent publications? Notably, the more heavily studied a gene was previously, the smaller the effect of GWAS association (Fig. 2B, Spearman rank correlation, *p*∼ 6 ×10^*-*8^, *N* = 2, 442).

The strength of a GWAS association is quantified by its statistical p-value and its biological effect size, which is most commonly an odds ratio. The normalized publication excess for a newly associated gene and the p-value of its association are weakly positively correlated (Fig. 2D; *p*∼ 1 ×10^*-*4^, *N* = 2, 442). GWAS results of greater statistical significance are thus correlated with greater additional publications. By contrast, the normalized publication excess is not significantly correlated with the effect size of the reported association (Fig. 2E; *p*∼ 0.16, *N* = 1, 295). Researchers are thus slightly more likely to follow up on associations with higher statistical confidence, rather than those with larger biological effects.

The strongest predictor of the effect of a GWAS on future publications for associated genes is, however, the year in which the GWAS was published. The normalized publication excess has declined dramatically since the early years of GWAS (Fig. 2F; *p*∼ 9 ×10^*-*23^, *N* = 2, 442).

The predictors for the effect of GWAS on subsequent publications that we have studied may themselves be correlated; to disentangle their effects, we built a linear regression model. In that model, the effects of the number of recent publications and the year of GWAS publication are strong and statistically significant (Table 1). By contrast, the quantitative properties of the association itself, the p-value and the effect size, are weak and not statistically significant.

**Table 1.**
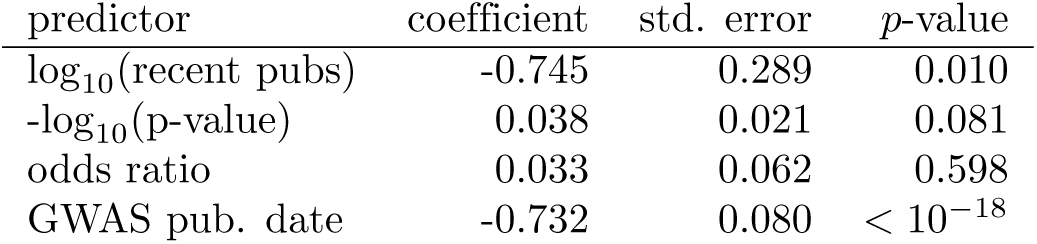
Linear regression model for the normalized publication excess of new GWAS genes (*N* = 1, 295).

Association with particular diseases might lead to particularly intense study. To test this possibility, we considered the class of disease that each gene was associated with as an additional predictor in the linear regression model. Of the 20 disease classes tested, only metabolic disease had a significant effect on the normalized publication excess (Table S1). Further stratifying among metabolic diseases, we found that this trend is driven by studies on type II diabetes and obesity (Table S2).

Association with complex disease via GWAS is correlated with an increase in subsequent publications on a gene (Fig. 2C). The magnitude of this increase does not depend strongly on the p-value or the effect size of the association, but it is smaller for genes that were more heavily studied or that were associated more recently (Table 1). The typical effect of GWAS on subsequent publications on an associated genes is thus declining, but does GWAS identify particular genes that receive exceptional study?

### Genes with exceptional publication records

The typical new GWAS gene experiences a modest increase in subsequent publications, but some exceptional genes may experience large increases, so-called “hot” genes. To identify such genes, we used the model of Pfeiffer and Hoffmann [25] to predict the number of publications for each gene in each year, based on that gene’s prior publication history. We trained the model on all genes never implicated in complex disease through GWAS. By comparing model predictions and publication data, we then identified particular years in which particular genes had unexpectedly large numbers of publications (Dataset S1). For example, Complement Factor H had a significant excess of publications in all three years following its association with macular degeneration (Fig. 3A).

**Fig 3.**
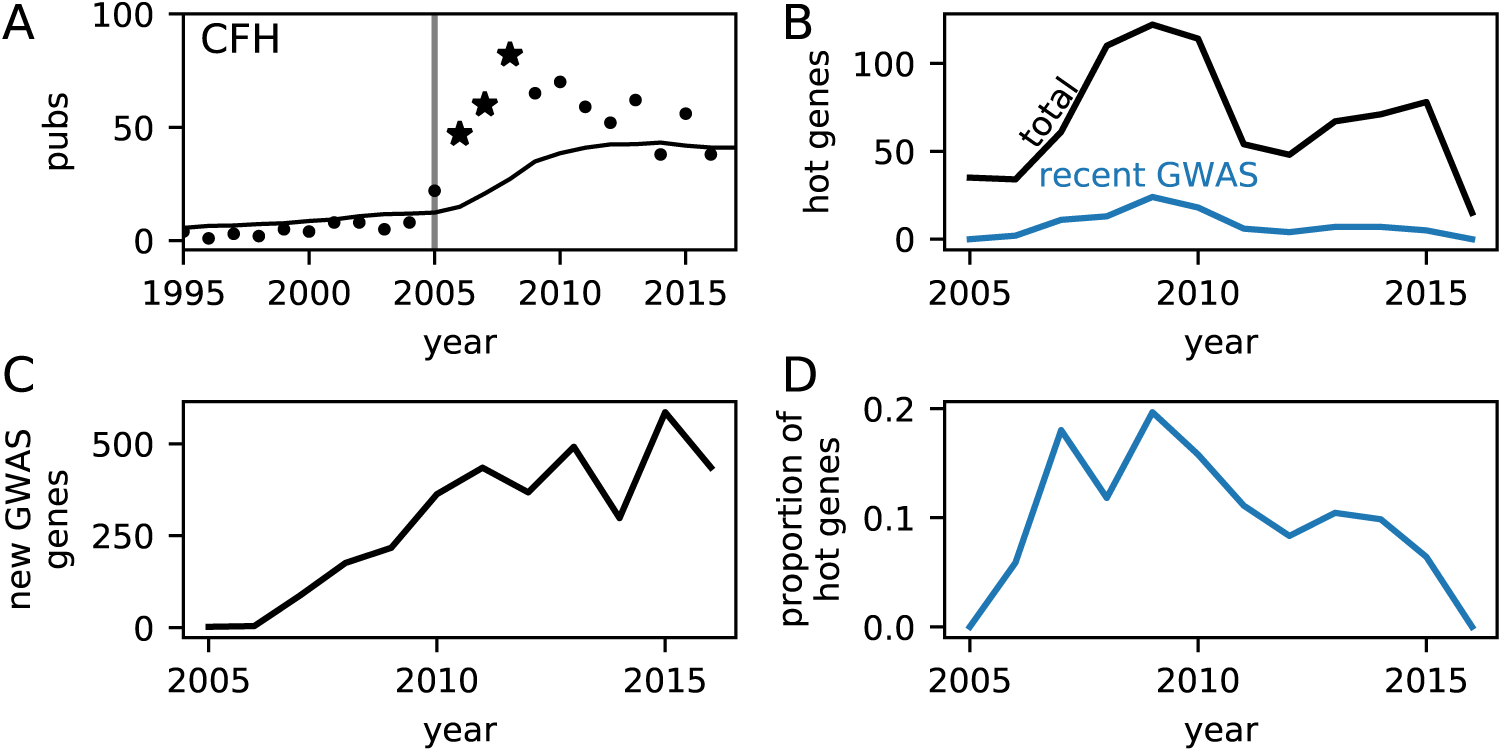
The effect of GWAS in generating exceptionally studied genes. A: A significantly elevated number of studies were published on Complement factor H following its association with macular degeneration via GWAS in 2005. Solid line is the predicted publication history from the model of [25], points indicate actual publication counts, and starred points indicate years with a statistically significant excess (one-sided Bonferroni-corrected *p <* 0.05). B: The total number of genes exhibiting an unusual excess in publications peaked in 2009, as did the number of those genes that were recently newly associated with complex disease via GWAS. C: The number of genes newly associated with complex disease through GWAS has grown since the inception of GWAS. D: The proportion of genes exhibiting an unusual excess in publications that were recently identified in GWAS peaked at roughly 20% in 2009 and has since declined.

The total number of hot genes per year has recently fluctuated (Fig. 3B). Between 2009 and 2016, on average 0.3% of genes were hot in any given year. Of genes that were newly associated with complex disease via GWAS within the past three years, the probability of being hot was 1.3%. So being newly associated with complex disease does increase the probability that a gene will become hot. The total number of hot genes that were recently associated with complex disease via GWAS peaked, however, in 2009 (Fig. 3B), even as the number of new GWAS genes each year has grown (Fig. 3C). Thus, the proportion of hot genes that were recent GWAS hits has declined markedly (Fig. 3D).

To further quantify the role of GWAS in creating hot genes, we used a logistic regression model (Table 2). Consistent with the overall probabilities (Fig. 3), this model showed that being a recent new GWAS hit was an important factor in determining whether a gene would be hot. The effect of being a GWAS hit, however, had a negative interaction with the year. In other words, the effect of GWAS on creating hot genes with exceptional publication records is decreasing with time.

**Table 2.** Logistic regression model for whether a gene exhibits a statistically significant excess in publications in a given year, compared to the expectation of the Pfeiffer and Hoffmann model [25].

| predictor | coefficient | std. error | p-value |
| --- | --- | --- | --- |
| $\log_{10}(\text{recent pubs})$ | 3.881 | 0.068 | $< 10^{-32}$ |
| year | -0.108 | 0.012 | $2 \times 10^{-18}$ |
| recent GWAS | 4.094 | 0.361 | $9 \times 10^{-30}$ |
| (year × recent GWAS) interaction | -0.569 | 0.064 | $1 \times 10^{-18}$ |

## Discussion

We analyzed biomedical research publications to quantify the effect of genome-wide association studies on published scientific research. We found that even after the advent of GWAS, publications remain highly skewed toward Mendelian disease genes, with many complex disease genes receiving little attention (Fig. 1C). New complex disease genes identified by GWAS do receive additional study and subsequent publications (Fig. 2C), but that effect has declined dramatically (Fig. 2F, Table 1). Being newly associated with complex disease does increase a gene’s chance of becoming a “hot” gene, but this effect has also declined (Fig. 3D, Table 2). Together, our results suggest that GWAS have been successful in bringing research attention to novel genes involved in complex human disease, but their influence is waning.

Considering the overall distribution of biomedical publications, we found that GWAS have not reduced the startling inequality among human genes. The distribution of publications among human genes is characterized by a Gini coefficient of 0.73 in the post-GWAS era (Fig. 1A). By comparison, the Gini coefficient of money income among American households was 0.48 in 2016 [26] and among global households was 0.625 in 2013 [27]. The inequality of publications among genes is thus substantially greater than the inequality of income among households.

Focusing on individual genes, we found that association with complex disease via GWAS is correlated with an increase in subsequent publications (Fig. 2). Interestingly, the p-value and effect size of the association play a statistically insignificant role in determining the magnitude of that increase (Table 1). Notably, the p-value is a somewhat better predictor, consistent with attention to statistical rather than biological significance [28]. We found a stronger effect on subsequent publications for genes newly associated with metabolic disease (Table S1 and S2), perhaps reflecting its recent emphasis in public health [29]. We also found that association with complex disease via GWAS does raise the chances of a gene becoming an exceptionally studied “hot” gene (Fig. 3). But most dramatically, we found that the effects of new association via GWAS have declined substantially over the past decade (Fig. 2F and 3D).

The direct results of a GWAS are associations of a disease with genetic variants, not with genes. For simplicity, we associated each variant with the closest gene, as long as that gene was within 500 kilobases. But many variants are regulatory, and gene regulation is complex, so some variants may actually most strongly affect other more distant genes [30]. Thus some of the gene associations we study may be spurious. But this issue has existed since the advent of GWAS and has not changed markedly since. So it cannot explain our most striking result, that the effect of GWAS on subsequent publications has declined over time. When studying the effects of genetic evidence on drug development, Nelson et al. [31] used a more complex approach for assigning variants to genes. They incorporated linkage disequilibrium and attempted to infer regulatory relationships using expression quantitative trait loci (eQTLs) and DNAse hypersensitivity sites. When we analyzed their collection of association data, we found similar results to our original analysis, although effects were somewhat weaker (Table S5 and Fig. S2). Notably, we still found a negative relationship between the publication date of an association and its effect on subsequent publications.

Our measures of scientific publications do not necessarily capture the full effects of GWAS on biomedical research. Motived by the example of Complement Factor H (Fig. 3A), we focused on publications in a three-year window following the GWAS. Some follow-up studies may take longer, but using a five-year window does not change our qualitative conclusions (Fig. S1 and Tables S3 and S4). GWAS may also promote biomedical research in ways that do not involve new publications. For example, drugs with associated genetic evidence are more likely to progress along the development pipeline [31], suggesting that GWAS promote efficient drug development. More broadly, we focused on associations with complex disease, the most common biomedical application of GWAS. But GWAS for drug response have already provided important guidance for personalized treatment [32]. Lastly, human GWAS have applications beyond health. For an evolutionary example, GWAS data have been used to detect adaptation in the human genome [33].

What explains our most striking result, the declining effect of GWAS on subsequent publications regarding newly associated genes? Perhaps early GWAS captured most genetic variants of large effect, so more recent studies find less compelling associations. But effect size is not a strong predictor of subsequent publications (Table 1). Moreover, the typical effect size of new associations has declined only modestly (Fig. S3A), and the absolute number of large-effect associations has grown (Fig. S3B). Or perhaps journal publication criteria have changed over time, making GWAS less visible or follow-up studies more challenging to publish. The typical impact factor of journals GWAS are published in has declined slightly since the advent of GWAS (Fig. S4A). But the impact factor of the GWAS publication has only a weak effect on the publication excess of newly associated genes (Fig. S4B). When we included GWAS publication impact factor in our linear regression model, its effect was statistically significant but insufficient to explain the effect of publication date (Table S6). Or perhaps the availability of funding for follow-up studies has declined, as overall biomedical research funding has declined in both North America and Europe [34]. Or perhaps the capacity and interest to perform follow-up analyses has not kept pace with the “fire hose” of GWAS results [35]. Our data do not point toward a definitive explanation, and further investigation is needed to understand why recent GWAS promote less follow-up study on associated genes than early GWAS.

Over the past decade, GWAS have undeniably contributed greatly to biomedical knowledge [11]. The development of large-scale accessible databases of phenotypic and genotypic data, such as the UK Biobank [36], will fuel further contributions. But few GWAS results are directly medically actionable, so follow-up research is essential to translate novel associations into medical innovations. Our results suggest that the ability of GWAS to motivate published follow-up research on associated genes is declining. To maximize the positive impact of GWAS on human health, this trend must be understood and reversed.

## Materials and Methods

### Publication data

We obtained Entrez GeneIDs for all 20,422 human protein-coding genes from NCBI Gene [23] on December 12, 2017. For all those genes, we collected PubMed identifiers of associated publications from NCBI Gene’s gene2pubmed file, downloaded December 12, 2017. This file contains both associations created manually during the curation of Gene References Into Function (GeneRIFs) and associations collected from organism-specific databases, Gene Ontology, and other curated data sources. We then obtained date information for each publication from PubMed, taking the earliest year between the reported Year or EYear, using BioPython [37]. We followed a similar procedure for yeast genes. We obtained impact factor data from the 2016 InCites Journal Citation Reports [38].

### Disease data

To identify genes associated with Mendelian disease, we downloaded the Online Mendelian Inheritance in Man (OMIM) Gene Map of connections from genes to traits [24] on January 17th, 2018. We filtered to keep only entries with a confidence code of “confirmed” and to ignore entries indicating a potentially spurious mapping or association with a non-disease trait. We further considered only entries with Entrez GeneIDs, to avoid ambiguity among gene names and aliases. This procedure yielded 1,878 genes associated with disease traits. Of these, 1,543 genes were associated with Mendelian but not complex multifactorial disease, 157 were associated with complex multifactorial but not Mendelian disease, and 178 were associated with both Mendelian and complex multifactorial disease.

To further identify genes associated with complex disease and to gather GWAS data, we used the January 1st, 2017 release of NHGRI-EBI’s GWAS Catalog [4]. We filtered the catalog to remove nondisease traits, by keeping only entries that were children of the term “disease” (EFO0000408) in the Experimental Factor Ontology [39]. To connect associated variants with genes, we began with the Mapped Genes column in the Catalog. We then connected each variant with its closest mapped gene, if that gene was within 500 kilobases. If a variant was within two overlapping genes, we connected with both genes. This procedure yielded 4,069 genes associated with complex disease. To analyze classes of disease, we used the children of the term “disease” in the Experimental Factor Ontology.

Our analysis of OMIM and the GWAS catalog yielded 5,369 total disease-associated genes. Considering genes associated with only Mendelian disease in OMIM and not associated with disease through GWAS yielded 1,126 Mendelian disease genes. Considering genes associated with only complex multifactorial disease in OMIM or associated with disease through GWAS yielded 3,648 complex disease genes. The remaining 595 genes we associated with both Mendelian and complex disease.

Of the disease genes in the GWAS catalog, 2,442 were first associated prior to 2015, so we could analyze three full years of publication data. For those genes, we identified odds ratios as effect sizes without units for variants that had a reported frequency of the risk allele. For our odds ratio analysis, we analyzed the 1,295 genes for which an odds ratio was reported in the first year of GWAS association.

We also analyzed the association data of Nelson et al. [31]. They connected variants to genes using linkage disequilibrium, expression QTLs, and DNAse hypersensitivity. We filtered their Supplementary Data Set 1 to remove associations from OMIM, which may be Mendelian diseases. We also manually classified traits as disease or non-disease (Dataset S2), filtering out the non-disease traits.

### Control genes

For each of our 2,442 GWAS genes, we identified its control gene as the non-GWAS gene with the closest number of total publications prior to the year the gene was first associated with complex disease. If multiple genes were tied for closest, we compared the previous year as well, continuing either until there was no ambiguity or until we reached 1950. For the 233 GWAS genes with ambiguous control genes, we compared subsequent publications between the GWAS gene and the average of the control genes.

### Publication rate model

We used the model of Pfeiffer and Hoffman to predict expected per-gene publication rates [25]:

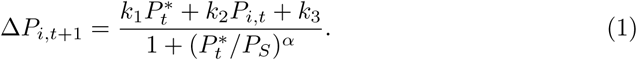

Here, Δ*P*_*i,t*+1_ is the predicted number of publications for gene *i* in year *t* + 1, and *P*_*i,t*_ and 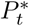 are the cumulative number of publications in previous years for the gene and the average cumulative number of publications for all genes in the organism, respectively. The term in the denominator models saturation of publication rates. The three rate parameters, *k*_1_, *k*_2_, and *k*_3_, and the saturation parameters, *P*_*S*_ and *α*, were assumed to be identical for all genes. To fit the parameters to our data, we constructed a likelihood function by assuming that the number of publications each year for each gene was independently Poisson distributed with mean Δ*P*_*i,t*+1_ given by Eq. 1. We then maximized that likelihood with respect to the five model parameters, using publication data from 1950 to 2015 for all non-GWAS genes. The maximum-likelihood parameter values were *k*_1_ = 0.0214, *k*_2_ = 0.225, *k*_3_ = 0.00288, *P*_*S*_ = 24.1, *α* = 1.67. Five genes each had 1 publication prior to 1950 that was not included in the data fit.

To identify years in which genes had significantly elevated publication rates, our null model was that publications were Poisson distributed with mean given by Eq. 1. Significant gene-years were defined as those in which the probability of generating at least the observed number of publications was less than the Bonferroni-corrected significance cutoff 0.05*/*(*N*_*g*_*N*_*y*_). Here *N*_*g*_ = 20, 442 was the total number of genes considered and *N*_*y*_ = 67 was the total number of years.

## Acknowledgments

We thank Yann Klimentidis and Tricia Serio for helpful comments.

---

**Table S1.** Effects of disease class on publication excess. We added parameters to the linear model designating whether or not a gene was first associated with a particular disease class. Only for metabolic disease do we see a statistically significant effect on publication excess. The second column shows the number of genes first associated with each disease class.

| predictor | # of genes | coefficient | std. error | p-value |
| --- | --- | --- | --- | --- |
| $\log_{10}(\text{recent pubs})$ | - | -0.739 | 0.292 | 0.012 |
| $-\log_{10}(\text{p-value})$ | - | 0.047 | 0.023 | 0.036 |
| odds ratio | - | 0.032 | 0.065 | 0.619 |
| GWAS pub. date | - | -0.760 | 0.085 | $10^{-18}$ |
| brain aneurysm | 4 | -1.950 | 2.877 | 0.498 |
| cardiovascular disease | 94 | -0.721 | 0.818 | 0.378 |
| chemotherapy-induced alopecia | 8 | 1.480 | 2.306 | 0.521 |
| digestive system disease | 138 | 0.431 | 0.703 | 0.540 |
| endocrine system disease | 10 | 2.825 | 1.969 | 0.152 |
| eye disease | 7 | 0.974 | 2.220 | 0.661 |
| genetic disorder | 40 | -0.611 | 1.085 | 0.573 |
| head and neck disorder | 20 | -0.087 | 1.371 | 0.949 |
| immune system disease | 317 | -0.881 | 0.536 | 0.101 |
| infectious disease | 34 | 0.282 | 1.141 | 0.805 |
| kidney disease | 29 | -1.500 | 1.220 | 0.219 |
| liver disease | 16 | -0.473 | 1.523 | 0.756 |
| mental or behavioural disorder | 132 | 0.031 | 0.735 | 0.966 |
| metabolic disease | 133 | 1.796 | 0.719 | **0.013** |
| neoplasm | 254 | -0.084 | 0.653 | 0.897 |
| nervous system disease | 221 | 0.436 | 0.632 | 0.490 |
| reproductive system disease | 27 | 1.046 | 1.216 | 0.390 |
| respiratory system disease | 41 | -0.436 | 1.035 | 0.674 |
| skeletal system disease | 73 | 1.090 | 0.845 | 0.198 |
| skin disease | 36 | -0.308 | 1.091 | 0.778 |

**Table S2.** Effects of different metabolic diseases on publication excess. We modified our linear model to remove the general metabolic disease parameter and add the three specific diseases with a large number of genes. Remaining model parameters (not shown) are similar to the regression in Table S1.

| predictor | # of genes | coefficient | std. error | p-value |
| --- | --- | --- | --- | --- |
| type I diabetes mellitus | 31 | -0.355 | 1.186 | 0.765 |
| type II diabetes mellitus | 60 | 6.271 | 0.922 | $10^{-11}$ |
| obesity | 31 | 6.249 | 1.162 | $10^{-7}$ |

**Table S3.** As in Table 1, linear regression model for the normalized publication excess of new GWAS genes (*N* = 873), but with a five-year range for calculating Δ pubs and recent pubs.

| predictor | coefficient | std. error | p-value |
| --- | --- | --- | --- |
| $\log_{10}(\text{recent pubs})$ | -0.558 | 0.524 | 0.287 |
| $-\log_{10}(\text{p-value})$ | 0.053 | 0.036 | 0.143 |
| odds ratio | -0.008 | 0.108 | 0.944 |
| GWAS pub. date | -1.554 | 0.191 | $10^{-15}$ |

**Table S4.**
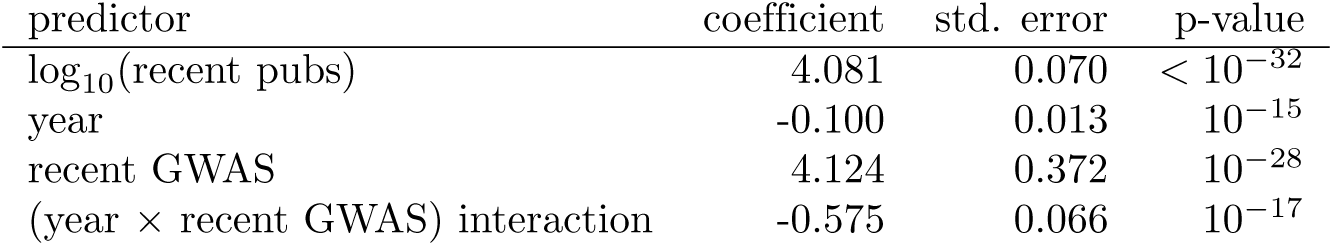
As in Table 2, logistic regression model for whether a gene exhibits a statistically significant excess in publications in a given year, compared to the expectation of the Pfeiffer and Hoffmann model [25], but with a 5-year range for defining recent pubs and GWAS.

**Table S5.** As in Table 1, linear regression model for the normalized publication excess of newly associated genes (*N* = 1, 861), but using the data of Nelson et al. [31]. Note that they did not collect effect size information. The negative effect of publication date remains, although it is somewhat weaker.

| predictor | coefficient | std. error | p-value |
| --- | --- | --- | --- |
| $\log_{10}(\text{recent pubs})$ | -0.119 | 0.235 | 0.614 |
| $-\log_{10}(\text{p-value})$ | -0.007 | 0.008 | 0.393 |
| GWAS pub. date | -0.448 | 0.073 | $10^{-9}$ |

**Table S6.** As in Table 1, linear regression model for the normalized publication excess of newly associated genes (*N* = 1, 191), but including the effect of impact factor for the GWAS publication.

| predictor | coefficient | std. error | p-value |
| --- | --- | --- | --- |
| $\log_{10}(\text{recent pubs})$ | -0.807 | 0.291 | 0.006 |
| $-\log_{10}(\text{p-value})$ | 0.026 | 0.022 | 0.235 |
| odds ratio | 0.050 | 0.062 | 0.420 |
| $-\log_{10}(\text{impact factor})$ | 0.422 | 0.192 | 0.028 |
| GWAS pub. date | -0.706 | 0.081 | $10^{-17}$ |

**Fig S1.**
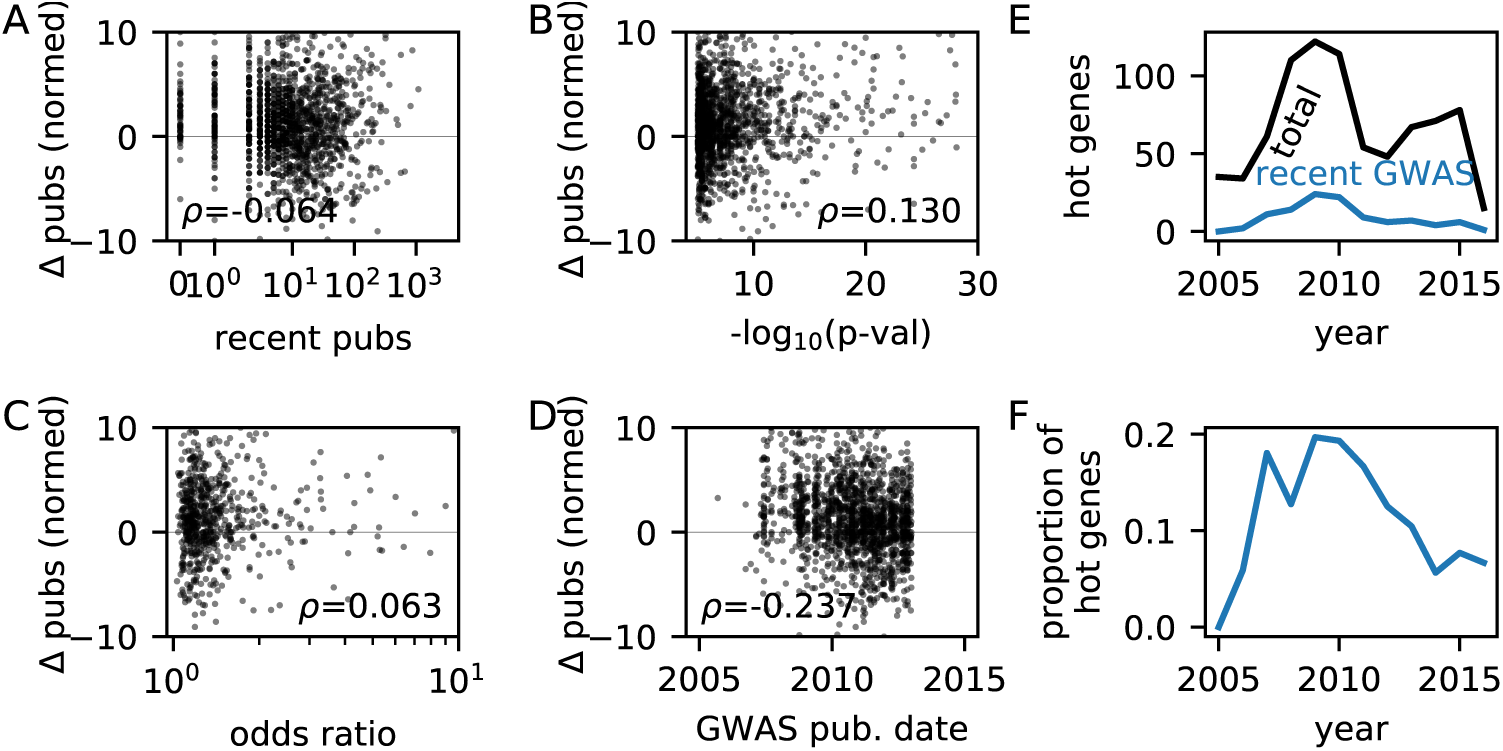
Results using a 5 year window for counting publications and defining recent GWAS. A-D: As in Fig. 2. E&F: As in Fig. 3.

**Fig S2.**
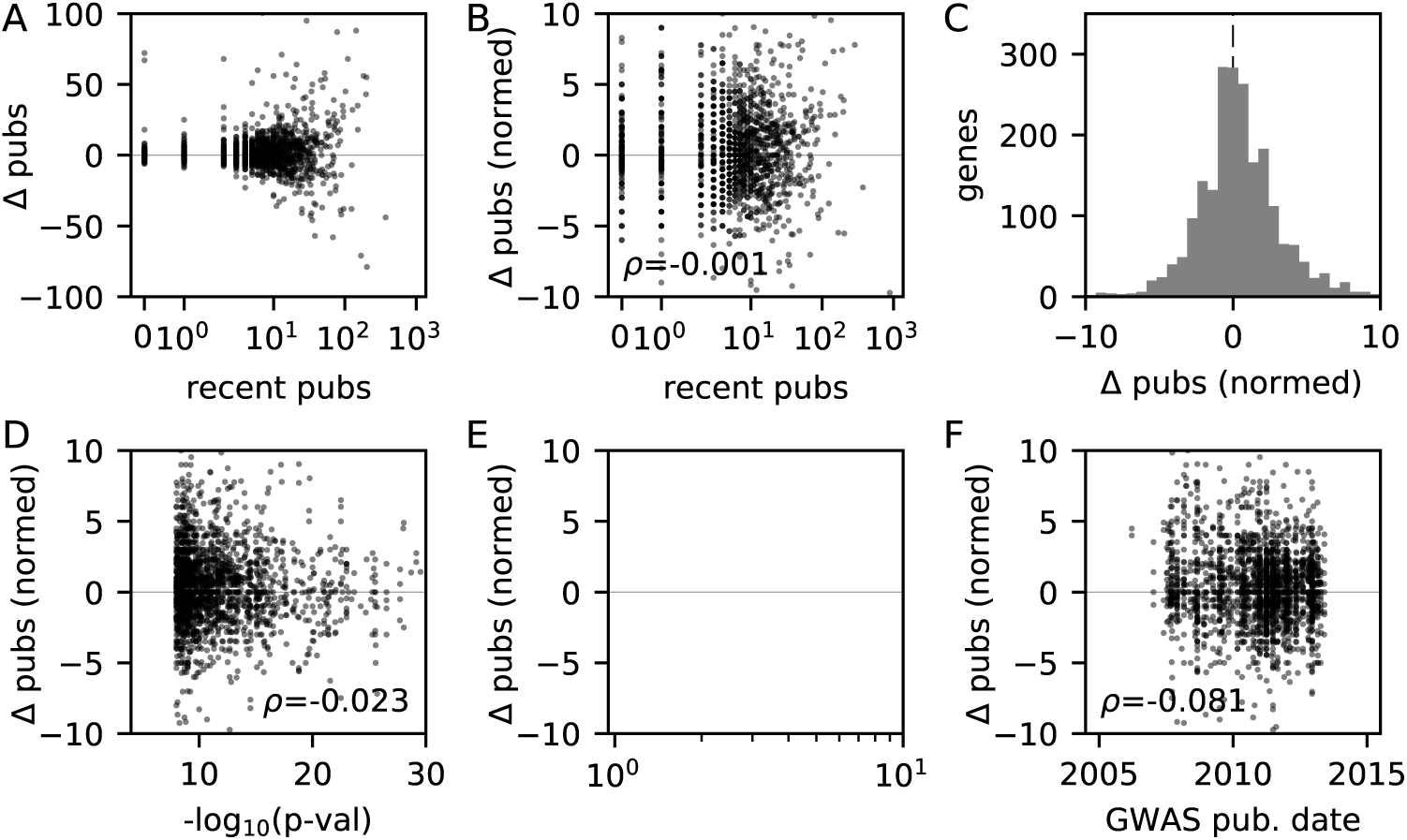
Results using association data from Nelson et al. [31] Panels as in Fig. 2, but panel F is empty because Nelson et al. did not collect odds ratios. Broadly, the effects are similar to those in our collection of data, but weaker. For the Nelson et al. data, the mean normalized publication excess is 0.71, compared to 1.24 in our collection of data.

**Fig S3.**
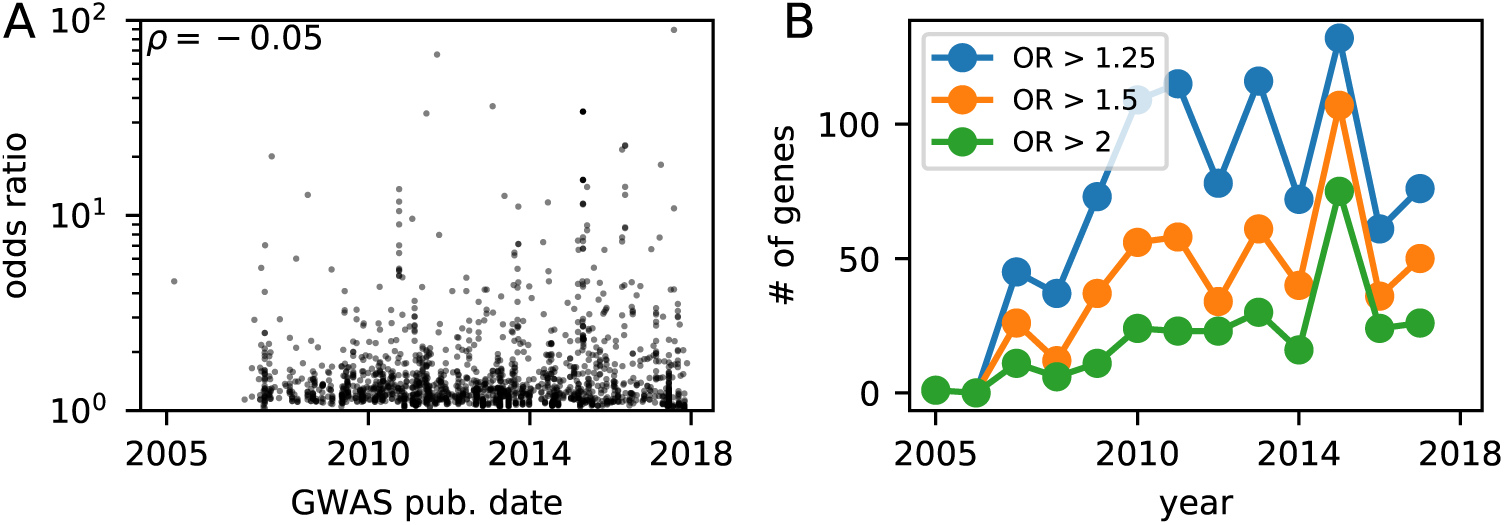
Trend in effect size of new GWAS genes. A: For genes newly associated with complex disease, the typical odds ratio has slightly declined over time (Spearman rank correlation *ρ* = −0.05, *p* ∼ 0.033, *N* = 1, 812). B: The number of new GWAS genes with large odds ratios has not, however, declined with time.

**Fig S4.**
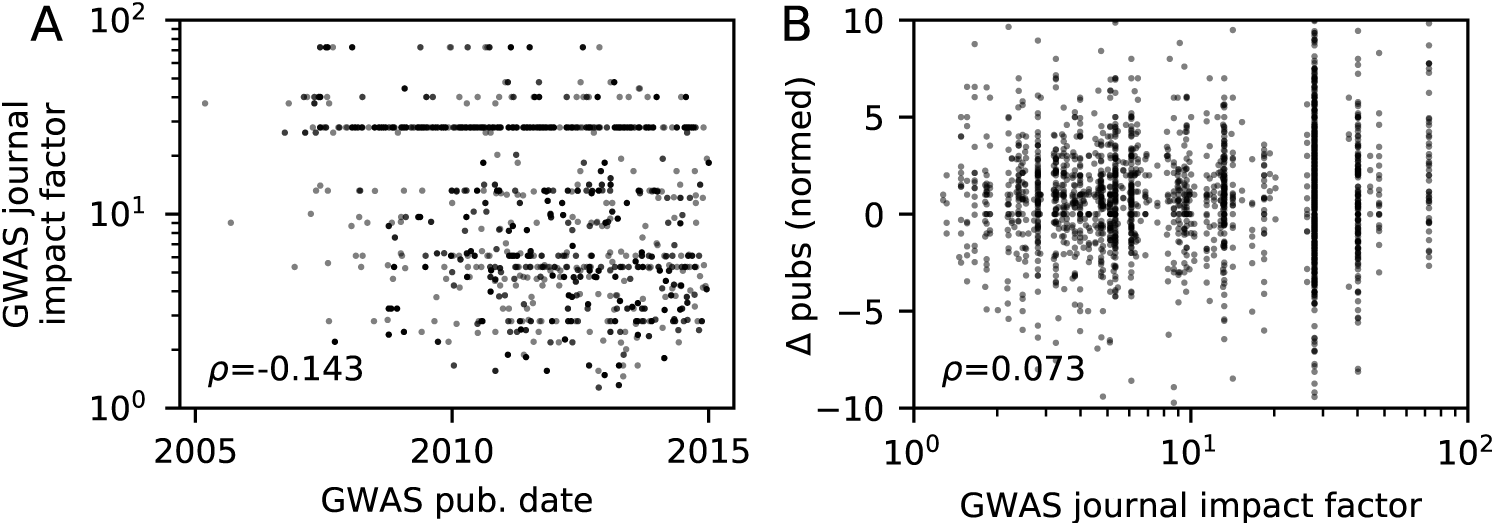
Effect of journal impact factor. A: The typical impact factor of GWAS publications has declined slightly since the advent of GWAS. B: The journal impact factor the GWAS publication is weakly correlated with publication excess of the reported associated genes.

**Dataset S1**: Gene-years with a statistically significant excess of publications relative to the prediction of the Pfeiffer and Hoffman model. For GWAS disease genes, the date of the first GWAS to identify that gene is also recorded.

**Dataset S2**: Traits from the association data of Nelson et al. [31], categorized as disease or non-disease.

